# Cell-type-specific methylome-wide association studies implicate neurodegenerative processes and neuroimmune communication in major depressive disorder

**DOI:** 10.1101/432088

**Authors:** Robin F. Chan, Gustavo Turecki, Andrey A. Shabalin, Jerry Guintivano, Min Zhao, Lin Y Xie, Gerard van Grootheest, Zachary A. Kaminsky, Brian Dean, Brenda W.J.H. Penninx, Karolina A. Aberg, Edwin J.C.G. van den Oord

## Abstract

We studied the methylome in three collections of human postmortem brain (N=206) and blood samples (N=1,132) of subjects with major depressive disorder (MDD) and controls. Using an epigenomic deconvolution approach we performed cell-type-specific methylome-wide association studies (MWAS) within sub-populations of neurons/glia and granulocytes/T-cells/B-cells/monocytes for bulk brain and blood data, respectively. Multiple MWAS findings in neurons/glia replicated across brain collections (ORs=509-538, P-values<1×10^−5^) and were reproducible in an array-based MWAS of sorted neurons/glia from a fourth brain collection (N=58). Pathway analyses implicated p75^NTR^/VEGF signaling, neurodegeneration, and blood-brain barrier perturbation. Cell-type-specific analysis in blood identified associations in CD14+ monocytes -- a cell type strongly linked to neuroimmune processes and stress. Top results in neurons/glia/bulk and monocytes were enriched for genes supported by GWAS for MDD (ORs=2.02-2.87, P-values=0.003 to <1×10^−5^), neurodegeneration and other psychiatric disorders. In summary, we identified novel MDD-methylation associations by using epigenomic deconvolution that provided important mechanistic insights for the disease.

---

Major depressive disorder (MDD) is a mental illness characterized by marked and persistent dysphoria^1^. Because the disease has high lifetime prevalence (~15%)^2^, can start early in life, and typically involves a chronic course, the World Health Organization ranks MDD as the leading cause of disability^3^. DNA methylation studies offer unique opportunities to better understand and treat MDD by improving our understanding of the involvement of DNA methylation in the dynamic features (e.g., episodic nature, course) of MDD and by providing insight into how environmental risks (e.g., stress) can impact symptom severity^4–6^. Importantly, methylation studies have profound translational potential, as methylation is modifiable by treatment and can potentially be used as biomarkers to improve diagnosis and clinical disease management.

Methylome-wide association studies (MWAS) are ideally performed in the tissues where the pathogenic processes likely manifest. There exists good evidence that MDD has a systemic component that involves both brain and peripheral immune cells^7, 8^. Therefore, we sought to characterize MDD-linked methylation changes in both brain and blood.

MWAS is typically performed using DNA from bulk tissues containing multiple cell types. Failure to account for these multiple cell types has several drawbacks^9^. Most recognized is the risk of false positive associations that occurs when the abundance of cell types varies across samples included in the study^10, 11^. Underappreciated is the negative impact of cell type heterogeneity on the statistical power to detect associations with disease. Thus, case-control differences may be of opposite directions between cell types, resulting in “diluted” and/or “canceled out” effect sizes in bulk tissue. Furthermore, as the most common cell types will drive the results, associations present in low abundance cells may remain undetected in bulk tissue. Finally, knowing what cell type harbors an association is key for the biological interpretation of findings and can be critical for designing proper functional follow-up experiments.

It is not practically or fiscally feasible to perform methylation assays on isolated cell populations at the sample sizes required for adequately powered MWAS. A practical solution is to apply statistical methods that are informed by data from reference sets of sorted cells to deconvolute the cell-type-specific effects from data from reference sets of sorted cells to deconvolute the cell-type-specific effects from data generated with bulk tissue^12^. This deconvolution approach is commonly used in expression studies but can readily be applied to methylation data^13, 14^. The method has been validated using pre-designed mixtures of cell types and applications using empirical data have confirmed its value by revealing associations undetectable in bulk tissue^15^.

In the largest and most comprehensive study to date, we examined methylation differences between MDD cases and controls in bulk brain samples from three collections totaling 206 individuals, as well as in 1,132 independent blood samples. Applying an epigenomic deconvolution strategy to bulk tissue data, we performed methylome-wide association studies (MWAS) within cell populations mainly consisting of neurons/glia and granulocytes/T-cells/B-cells/monocytes. Associations detected in one set of brain collections were replicated in the others using a stringent “round-robin” design. We further validated top cell-type-specific associations obtained via epigenomic deconvolution against those observed in sorted neuronal and glial nuclei from brains of an additional 58 case-control subjects. Finally, we tested for overrepresentation of genes implicated by our top MWAS findings among those identified in recent GWAS for MDD and related disorders.

## RESULTS

Complete descriptions of study participants, data quality control, and analyses are provided in the **Supplementary Methods**.

### Cell-type-specific MWAS in brain

We used sequencing-based methylation data from a total of 206 postmortem brain samples from three collections^16^. The sample collections were predominantly from Australia (AUS; 30 MDD, 31 control; Brodmann Area [BA] 25), United States of America (USA; 44 MDD, 37 control; BA10) and Canada (CAN; 39 MDD, 25 control; BA10). Overall, brain sample characteristics for MDD cases and controls were similar (**Table S1**).

To obtain sequencing-based reference methylomes for neurons and glia, we used fluorescence-activated cell sorting (FACS) to isolate neuronal and glial nuclei from cortex of five individuals (**Online Methods**). These reference methylomes enabled us to test for cell-type-specific case-control differences in samples for which only bulk tissue data was available. The principles underlying this epigenetic deconvolution approach^13, 15^ are illustrated in **Figure 1**. Implementation details for the method are presented in **Online Methods** and **Supplemental Note 1**.

**Figure 1:**
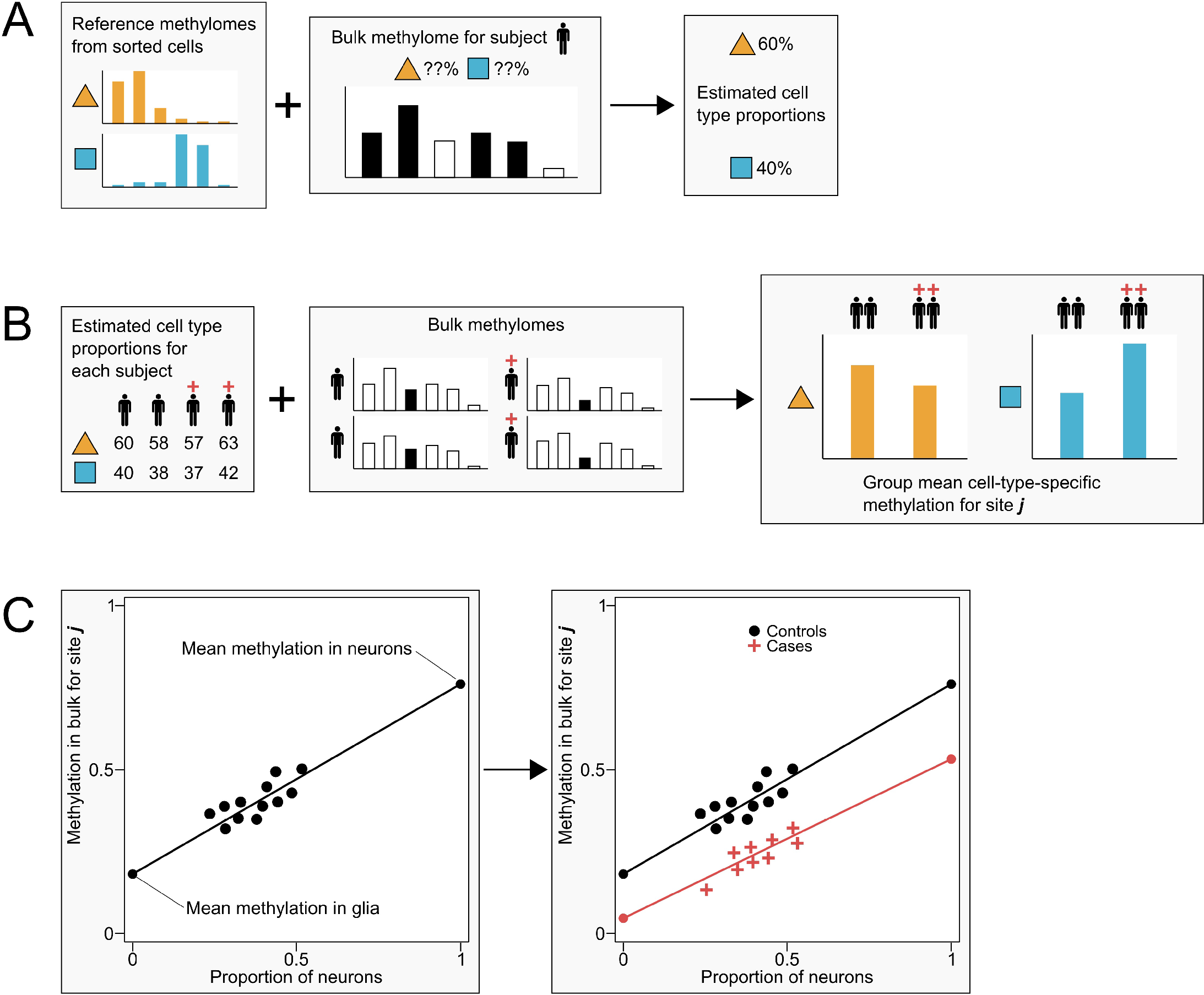
Epigenomic deconvolution for testing case-control differences in subpopulations of cells. **(A)** In step 1 of the deconvolution method, reference methylomes from purified samples of sorted cells are used to estimate cell type proportions. For each subject at a time, bulk methylation data is regressed on the most informative sites in the reference methylomes to obtain estimated proportions of each cell type. **(B)** Once cell type proportions have been estimated for each subject in the study, step 2 uses these proportions to estimate case-control differences at site each CpG site at a time. **(C)** To illustrate how cell-type-specific differences are estimated we present a simple example. Since bulk methylation and proportions of neurons/glia will differ between subjects, we can regress bulk methylation levels (Y-axis) on the proportion of neuronal cells (X-axis). Thus, extrapolating the regression line to the point where the proportion of neurons is zero (i.e., there are only glia cells) estimates the group mean methylation in glia, and extrapolation to the point where the proportion of neurons is one estimates the group mean methylation in neurons. By allowing the regression lines to differ between controls (black dots) and cases (red crosses), we obtain different predicted cell-type-specific group means that can be tested for significance using standard statistical tests. See **Online Methods** and **Supplemental Note 1** for discussion of the statistical models.

Estimated proportions of neurons and glia (~1:3) matched proportions expected in cortex based on the literature^17^ and showed no significant case-control differences. Quantile-Quantile (QQ) plots for the cell-type-specific MWASs (**Figure S3**) displayed deviations from the 95% confidence interval for small P-values suggesting multiple CpGs had discernible effects within cell types. Further, MWASs of permuted case-control status for each analysis yielded average lambdas that were not significantly different from 1 (**Figure S4**), which indicated that our observed P-values were accurate and did not show evidence of inflation. Additional validation analyses for the deconvolution method are detailed in the **Online Methods**.

To replicate findings in brain we used a stringent round-robin design (**Figure 2**). In each round, a meta-analysis of two of three datasets (“mini-meta”) was used for discovery (P<0.01), and the remaining dataset was used for replication (P<0.05), where direction of effect for discovery/replication must be equal. Markers that met these criteria in at least two of the three round-robin iterations were considered to have replicated.

**Figure 2:**
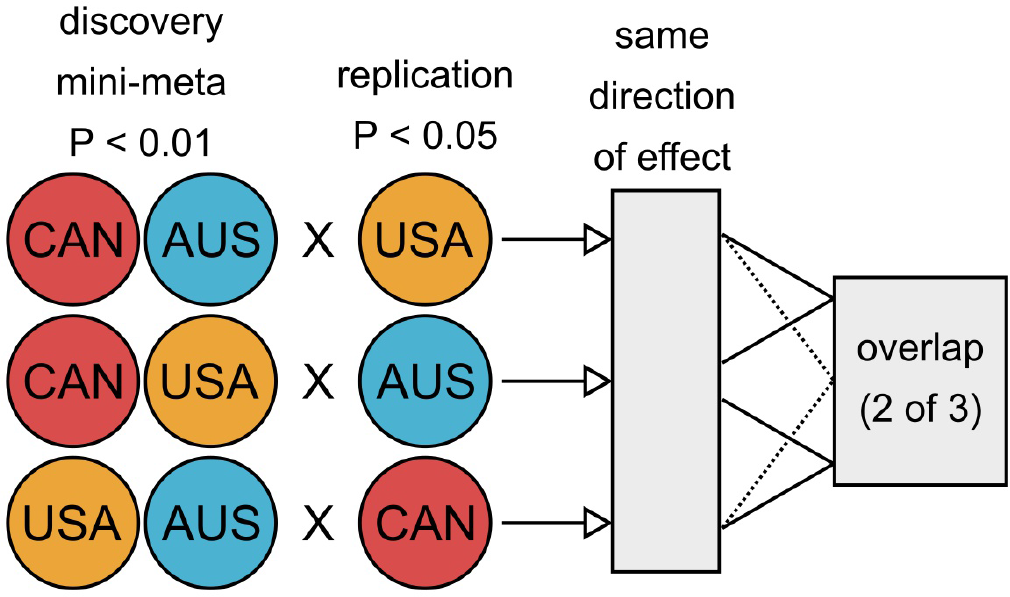
Round-robin replication design for neuron/glia/bulk MWAS in three brain collections. We performed a discovery MWAS by performing a meta-analysis using only two of the three individual MWASs. Results from this discovery “mini-meta” were then replicated in the remaining independent. Within each of the three possible rounds, a P-value threshold of 0.01 was used for discovery mini-meta results. Top discovery sites were considered to replicate if P<0.05 in the replication set and filtered for equal direction of effect. Finally, only replicating sites that were implicated in at least two of the three possible round-robin iterations were considered to have survived the protocol.

Permutation tests (**Table 1**) indicated that the replication of markers across datasets was not due to chance for the neuronal (odds ratio [OR]=509, P<1×10^−5^), glial (OR=518, P<1×10^−5^), or bulk (OR=538, P<1×10^−5^) analyses.

**Table 1.**
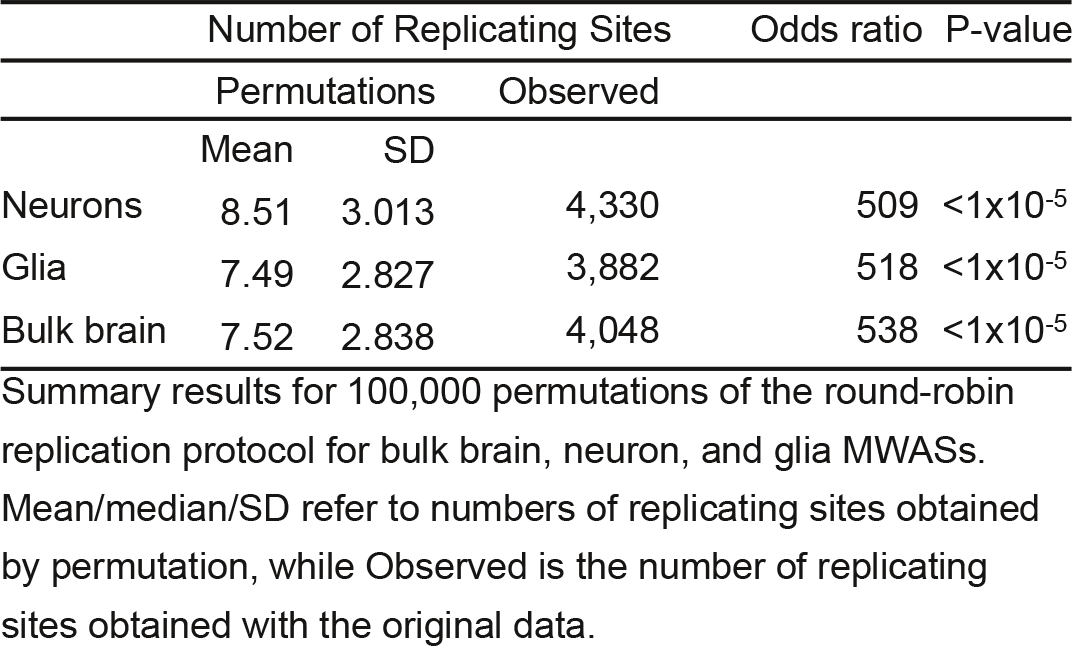
Sites passing round-robin replication

#### Neurons & Glia

A total of 4,330/3,882 replicating CpGs that implicated 1,784/1,682 genes were observed for neurons and glia, respectively (**Tables S2 & S3**). Interestingly, 1,683 CpGs were detected in both neurons and glia and were found in genes such as *HERC2*, *RNF111*, and *TRIM3* that all encode ubiquitin ligases that have roles in neurodevelopment and synaptic plasticity^18–23^. *HERC2* has also been previously associated with autism^24^. Top findings that were unique to neurons included genes like *RAPGEF6* and *FAM63B*. Interestingly, *RAPGEF6* has also been strongly associated with schizophrenia and has been shown to impact anxiety-like behavior in mice^25, 26^. Encoding the deubiquitinase MINDY2, *FAM63B* was a top finding from two past methylation studies of schizophrenia^27^ and bipolar disorder^28^. Finally, among top unique glia findings was a site within the gene *HMCN1*, which has been previously associated with post-partum depression^29^.

Considering the known effects of DNA methylation on distal regulatory elements, we tested whether our MWAS findings overlapped with Roadmap Epigenomics Project chromatin state tracks^30^. In general, results for neurons and glia were not overrepresented at chromatin states associated with regulatory features (**Table S4**). However, as the Roadmap Epigenomics Project chromatin state tracks for brain were generated in bulk tissue, they likely are not representative for many cell-type-specific chromatin states.

We tested results for enrichment of KEGG/ Reactome pathways using a permutation-based method that properly controls for the number of CpGs in a gene and the presence of correlated sites. Furthermore, this approach allows for a correction for testing multiple pathways with overlapping genes (**Online Methods**). To identify groups of pathways driven by the same MWAS findings, we further clustered the pathways that remained significant after correcting for multiple testing (family-wise error rate<0.05) based on the presence of overlapping genes.

Results for neurons were significantly overrepresented in genes belonging to 14 pathways (**Table S5**) and formed five clusters (**Figure 3A**). The first cluster (red) was related to *p75 NTR receptor-mediated signaling* (OR=3.84, P=0.0017) and related neuronal cell death processes. A second cluster (yellow) was related to *Glutamatergic synapse* (OR=3.97, P= 0.0123) and *Circadian entrainment* (OR= 4.47, P=0.0023). The most significant pathways in the remaining clusters were *Adherens junction* (OR=4.77, P= 0.0031), *Pathways in cancer* (OR=1.69, P=0.0033), and *Rho GTPase cycle* (OR=3.80, P=0.0042).

**Figure 3:**
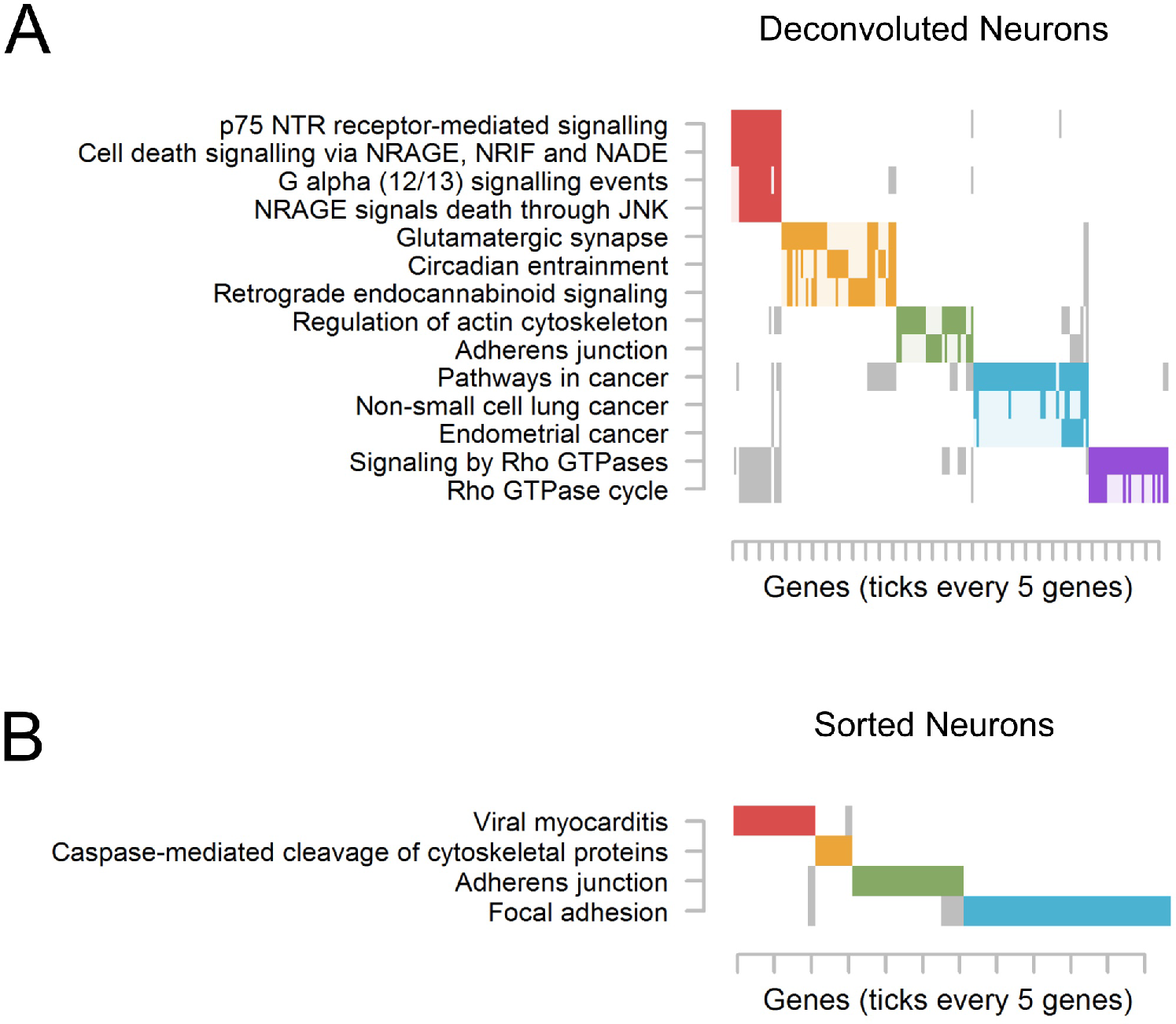
Cluster plot of significantly enriched pathways for neurons. As pathways often share genes, the raster plot visualizes the clustering of pathways (y-axis) determined on the basis of their overlapping genes (x-axis). The solid rectangles indicate genes that were both among the top MWAS results and a member of the listed pathway. Note, that only genes that were among the top MWAS results are plotted, rather than all possible pathway members. Only pathways containing a minimum of 4 overlapping genes and those passing family-wise significance were retained. Complete pathway names, gene names, odds ratios, and P-values are presented in **Table S5** for deconvoluted neurons and **Table S9** for sorted neurons.

Genes implicated by results in glia were overrepresented for 14 pathways (**Table S6**) that segregated into eight clusters (**Figure 4A**). The largest cluster (red) of glial pathways was driven by multiple classes of related secondary messenger systems such as *Ca-dependent events* (OR=6.53, P=0.0056) and *PLC beta mediated events* (OR=4.60, P= 0.0190). Other notable pathways enriched in results for glia were *p75 NTR receptor-mediated signaling* (OR=3.48, P=0.0095), *Role of ABL in ROBO-SLIT signaling* (OR=13.7, P=0.0282), and *Cortisol synthesis and secretion* (OR=3.73, P=0.0484).

**Figure 4:**
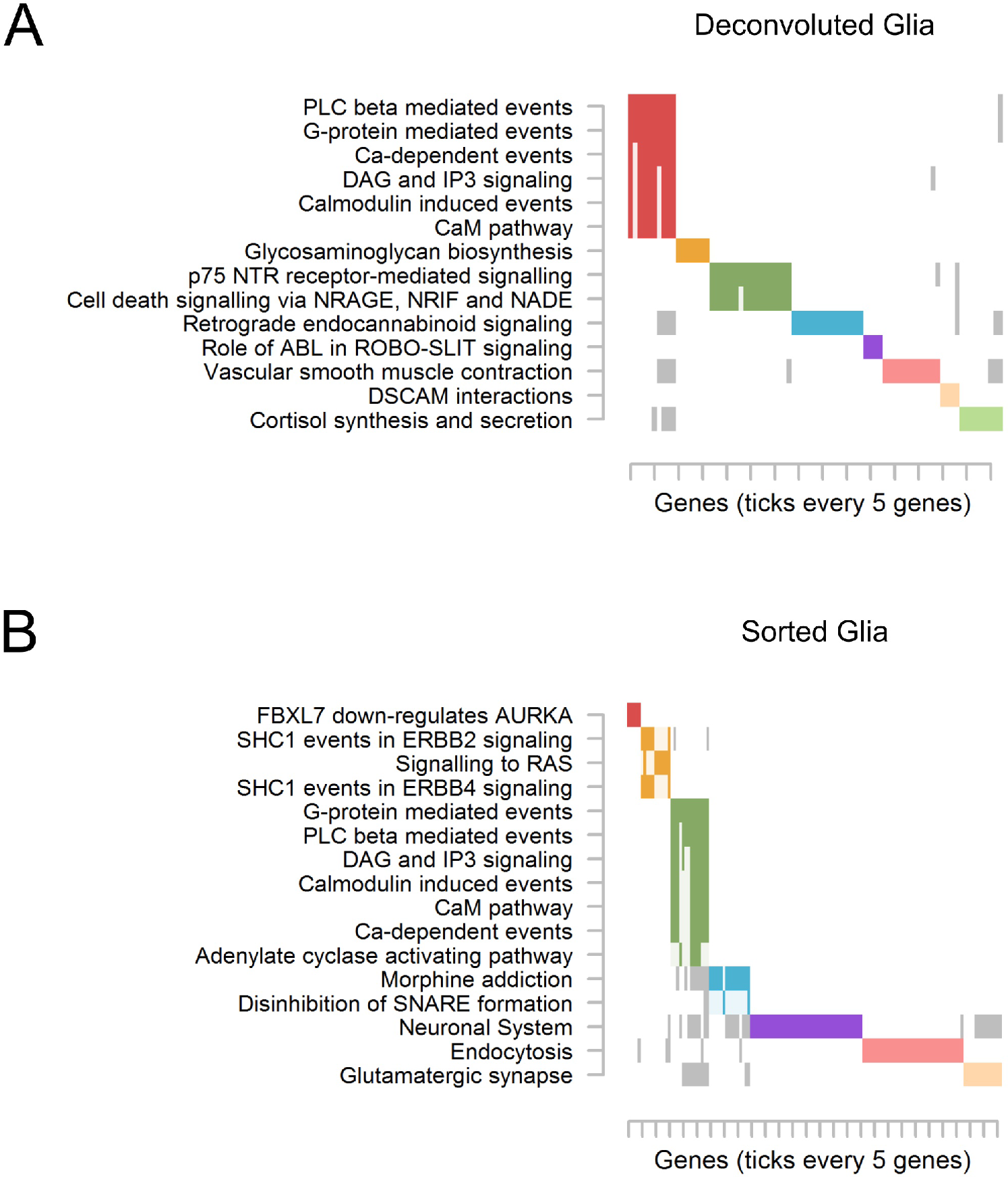
Cluster plot of significantly enriched pathways for glia. See **Figure 3** for an explanation of the plot. Complete pathway names, gene names, odds ratios, and P-values are presented in **Table S6** for deconvoluted glia and **Table S10** for sorted glia.

#### Complementary Analyses in Sorted Neurons & Glia

To check the cell-type-specific results obtained via epigenomic deconvolution, we also performed MWAS on array-based methylation data from FACS isolated neuronal (28 MDD, 29 control) and glial (29 MDD, 29 control) nuclei of postmortem frontal cortex samples31 (**Figures S7 & S8**). We next tested for enrichment between the replicating sites from our sequencing-based cell-type-specific MWASs (above) and the top results from the array-based sorted datasets (**Tables S7 & S8**), using two P-value thresholds of 0.05 and 0.01 for replication in the latter. It should be noted that only a small fraction of CpGs assayed in our sequencing-based data are also assayed by array-based approaches. For example, of the 4,330 replicating CpGs for deconvoluted neurons, only 169 could be mapped to CpGs actually measured in the array-based dataset.

Nonetheless, despite the limited sample size and scope of the sorted data, results for deconvoluted neurons were significantly enriched among top results for sorted neurons (19 CpGs, OR=2.05, P=0.0484). Our results for deconvoluted glia were also enriched among top (P<0.01) results for sorted glia (7 CpGs, OR=3.11), but did not remain significant after correcting for multiple thresholds (P=0.0661).

Top results (P<0.01) from MWAS of FACS sorted neurons and glia were significantly overrepresented for a number of pathways (**Tables S9 & S10**) that were also implicated in deconvoluted neurons and glia. Notably, sorted neurons (**Figure 3B**) were enriched for *Caspase-mediated cleavage of cytoskeletal proteins* (OR=7.67, P=0.0165) which is a central apoptotic process and involved in neuronal cell death^32, 33^. Pathways involving adherens junctions and focal adhesion molecules were similarly shared between results for sorted and deconvoluted neurons. Remarkably, the major pathway cluster (green, **Figure S4B**) for sorted glia contained terms related to, for example, *Ca-dependent events* (OR=4.76, P=0.0074), which closely mirrored results for deconvoluted glia. Together, these results strongly support the veracity and robustness of the cell-type-specific effects detected by the epigenomic deconvolution approach.

#### Bulk Brain

Analysis of bulk brain may provide better power to detect case-control differences that influence multiple cell types in a similar fashion. Therefore, we also applied the round-robin protocol to MWAS of the bulk brain methylation data, identifying 4,048 MDD-associated CpGs that replicated and implicated 1,786 genes (**Table S11**). The top findings in bulk brain included sites located in *RBFOX1*, *TMEM44*, and *PREX1*. Variants in the RNA-slicing regulator *RBFOX1* obtained genome-wide significance in a recent meta-analysis of large MDD GWASs^34^. In plasma, *TMEM44* has been associated with circulating levels of the pro-apoptotic tumor necrosis factor receptor 1 (TNFR1)^35^. In rodent models, deficits in *Prex1* results in autism-like behavior^36^ and is associated with anti-depressant response in humans^37^. Interestingly, very few associations that were detected in deconvoluted neurons (1 CpG) or glia (14 CpGs) were also detected in bulk, suggesting cell-type-specific effects are indeed diluted or obscured in bulk tissue.

Results for bulk brain were overrepresented in regulatory regions such as *Bivalent/Poised TSS* (OR=2.77, P=0.0003) and *Flanking Bivalent TSS/ Enhancer* (OR=2.70, P=0.0238) (**Table S4**). A total of 10 pathways (**Table S12**) were significantly overrepresented among the results for bulk brain spread across five clusters (**Figure 5**). Processes related to *Signaling by VEGF* (OR=4.03, P=0.0015) and *Nitric oxide stimulates guanylate cyclase* (OR=8.78, P=0.0054) were representative for the first (red) and second (yellow) clusters. Other notable pathways included *Focal adhesion* (OR=2.60, P=0.0424) and *NCAM signaling for neurite out-growth* (OR=4.61, P=0.0095).

**Figure 5:**
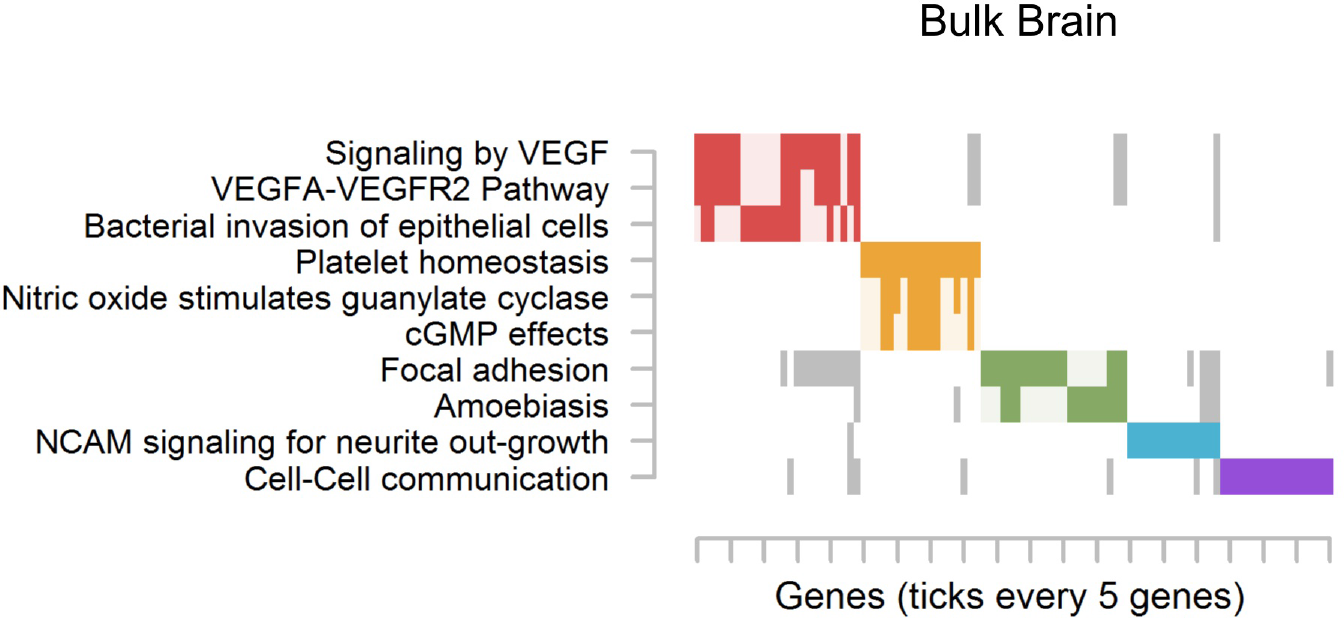
Cluster plot of significantly enriched pathways for bulk brain. See **Figure 3** for an explanation of the plot. Complete pathway names, gene names, odds ratios, and P-values are presented in **Table S12** for bulk brain.

### Cell-type-specific MWAS in blood

For blood, we used sequencing-based data from 1,132 whole blood samples^16^ (812 MDD and 320 controls) from the Netherlands Study of Depression and Anxiety (NESDA)^38^. For deconvolution of blood cell types, we used sequencing-based references^39^ from leukocyte populations isolated from six human whole blood samples using antibodies against CD15, CD3, CD19, and CD14 that are expressed on the surface of granulocytes, T-cells, B-cells, and monocytes, respectively^40^.

Mean estimated cell type proportions were 55.8, 31.3, 9.3, and 3.6% for CD15, CD3, CD19, and CD14 respectively. With the exception of CD14, the estimated cell type proportions differed significantly between cases and controls. Notably, MDD cases tended to have increased myeloid cell and decreased lymphocyte levels as expected^41, 42^.

#### CD15, CD3, CD14, & CD19

As a complementary dataset was unavailable for replication of the NESDA sample, we applied an appropriate false discovery rate (FDR) of 0.1 for declaration of methylome-wide significance^43, 44^ in cell-type-specific MWAS in blood. The QQ-plots (**Figure S5**) suggested the main association signals involved CD3 (T-cells) and CD14 (monocytes), which yielded multiple methylome-wide significant results. No significant findings from cell-type-specific MWAS in CD15 or CD19 were observed. Permutations of case-control status for each MWAS in CD3 and CD14 displayed mean lambda values that were not significantly different from 1 (**Figure S6**), again suggesting the observed effects were not due to uncontrolled artifacts.

In CD3, 18 CpGs passed methylome-wide significance (**Table S14**). Genic findings for CD3 involved *STRADB*, *FLI1/SENCR*, and *KIAA1217*. Due to the scarcity of methylome-wide significant results for CD3, functional annotation and pathway analyses are not presented.

The MWAS for CD14 identified 372 methylome-wide significant CpGs representing 129 genes (**Table S15**). Among the top genic findings for CD14 were *ITPR2*, *SVOPL*, *TP53*, *ARNT2*, *SHANK2*, *KATNAL2*, and *GRIA1*. Findings for CD14 were significantly enriched (OR=52.8, P=0.0063) at active transcriptional start sites for monocytes (**Table S4**). Top CD14 MWAS results showed overrepresentation of genes involved in 15 pathways (**Table S16**) that resulted in five clusters (**Figure 6**). The largest pathway cluster in CD14 (red) contained pathways involving *Glutamatergic synapse* (OR=15.0, P=0.0007) and *Oxytocin signaling pathway* (8.85, P=0.0061).

**Figure 6:**
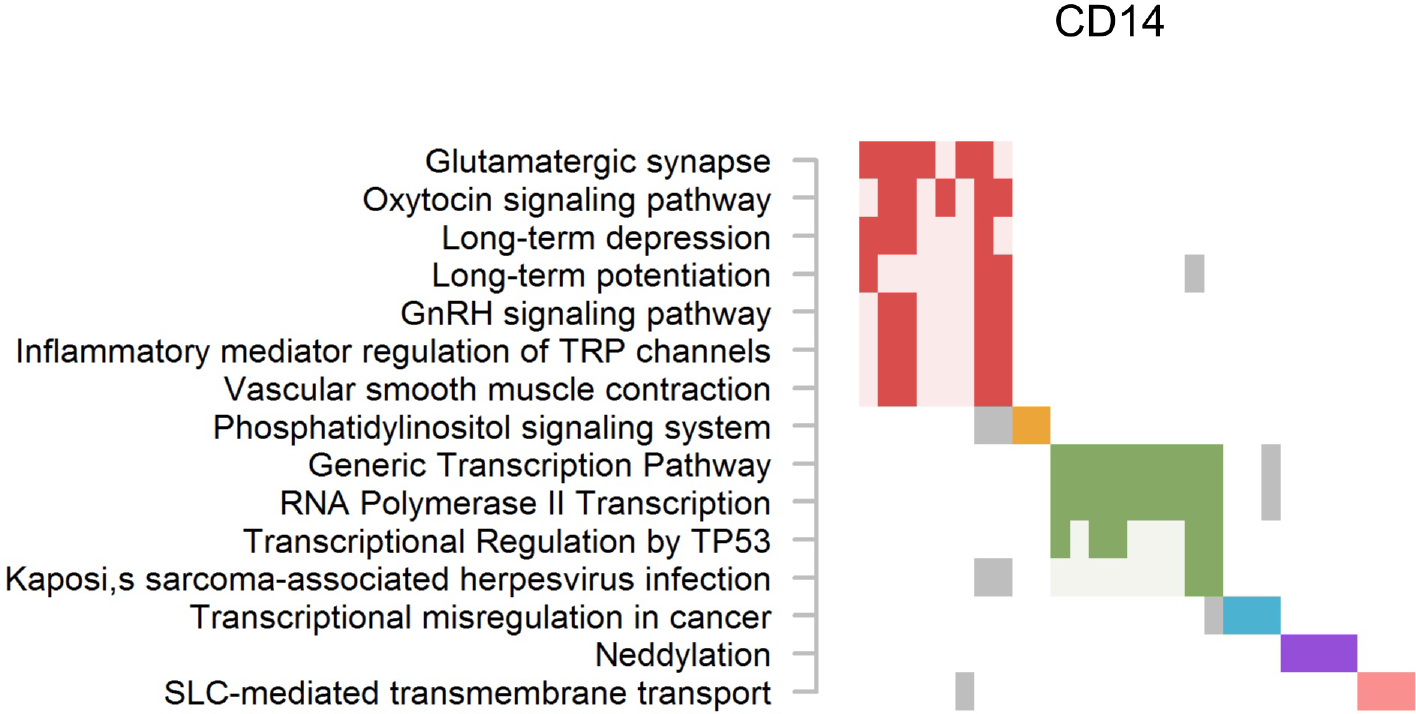
Cluster plot of significantly enriched pathways for CD14. See **Figure 3** for an explanation of the plot. Complete pathway names, gene names, odds ratios, and P-values are presented in **Table S16** for CD14.

#### Whole Blood

Top findings in whole blood^16^ (e.g., top site P=1.91×10^−8^) did not pass the FDR threshold of 0.1 employed in the current analysis. This again suggests that many effects in individual cell types may counter each other and leave many associations obscured in whole blood. Alternatively, as the blood cell types showing the largest signals are of relatively low abundance (CD3 and CD14), statistical power may be lacking to detect many of these differences as they represent a minority of cells.

### Cell-type-specific MWAS of antidepressant treatment

Given the potential for antidepressant drug treatment to affect the methylome^45^ we also sought to determine the cell-type-specific effects of drug treatment. Biographical information for postmortem brain samples often lack treatment history and precluded such an analysis in the brain datasets. However such information was available in NESDA, where we performed cell-type-specific MWAS with MDD cases that were treated (N=450) or untreated (N=362) with antidepressants. Antidepressant treatment was associated (FDR=0.1) with 3 CpGs in CD3 and 359 CpGs in CD14. Importantly, no sites that were associated with antidepressant treatment were also among top MDD findings in CD3 and CD14 MWASs. Thus, cell-type-specific associations to MDD in CD3 and CD14 were not simply due to drug treatment, and vice versa.

### Top findings are overrepresented at genes from GWAS of MDD and other neuropsychiatric disorders

We looked for convergence of evidence between our cell-type-specific MWASs and the top 10,000 variants from six recent GWAS meta-analyses for attention-deficit/ hyperactivity disorder (ADHD)^46^, anxiety disorders^47^, autism spectrum disorder (ASD)^48^, bipolar disorder (BPD)^49^, MDD^34^, and schizophrenia^49^. Additionally, given that our results consistently implicated pathways involved in neuronal apoptosis, we looked for overlap between MWAS results and 869 top GWAS sites for neurodegenerative disorders from NHGRI-EBI GWAS Catalog data^50^.

Genetic variants and methylation markers likely exert effects on a given gene at distally remote loci (e.g. promoters versus distal protein-coding sequence).
Therefore, we tested for significant enrichment between the genes implicated by top MWAS and GWAS sites, while accounting for local correlations and number of sites per gene (**Online Methods**). To check for specificity, we also tested for overlap of our top MWAS findings versus the top 10,000 variants from a recent GWAS meta-analysis of breast cancer^51^.

As expected, no MWAS results were enriched at genes associated with breast cancer. In contrast, results (**Table 2**) showed very robust and highly significant enrichment between genes implicated by MDD GWAS and those replicating across neuron, glia, and bulk brain MWAS results. Albeit less robust, replicating CD14 MWAS results were also significantly enriched at genes from MDD GWAS. Testing also yielded significant overlap of genes between bulk brain MWAS and GWAS for BPD and neurodegenerative disorders. Results for neuron MWAS were further enriched for genes associated with ADHD, ASD, and BPD. Glial MWAS findings were also enriched for ADHD and BPD genes, as well as those for neurodegenerative disorders.

**Table 2.**
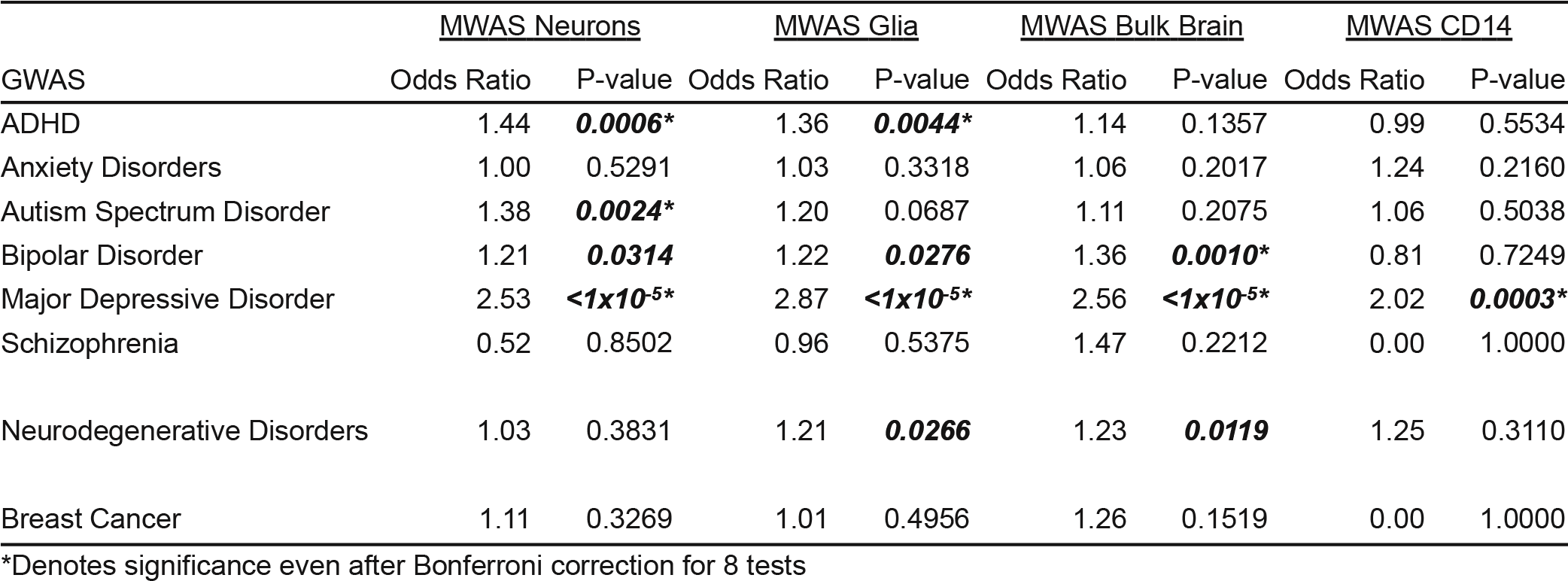
Enrichment of top MWAS findings versus GWAS of neuropsychiatric disorders

## DISCUSSION

In the most comprehensive methylation study of MDD to date, we characterized methylome-wide associations in large collections of brain and blood samples at a cell-type-specific level. Using a round-robin replication procedure, we identified novel associations with MDD in neurons and glia that replicated across three brain collections and in a fourth sample of sorted nuclei. Cell type-specific MWASs in blood uncovered associations in CD14+ monocytes that were not detected in whole blood. Strong overlap with past GWAS studies of MDD and neurodegenerative disorders also supported the robustness of our MWAS findings.

We obtained our results by employing an epigenomic deconvolution strategy to perform MWAS on individual sub-populations of neurons/glia and granulocytes/T-cells/B-cells/monocytes, respectively. The robustness of this strategy was demonstrated through a series of validation analyses (**Online Methods**). Cell-type-specific associations for deconvoluted neurons/glia also replicated across brain sample collections. Critically, overlap in terms of genes and pathways was found with MWASs that involved sorted neurons/glia. Finally, top results from cell-type-specific MWASs were significantly enriched for genes implicated by external GWAS of MDD and related disorders. Taken together, this converge of evidence supports the value of the deconvolution strategy as a cost-effective approach to detect cell-type-specific associations.

The overall results across brain cell types heavily implicated neurotrophin-linked degenerative pathways in MDD. The nerve growth factor receptor (p75^NTR^) regulates neuronal apoptosis via interacting proteins such as NRAGE, JNK, and Rac. A fine balance of neurotrophin signaling through the generally pro-survival Trk receptors and apoptotic p75^NTR^ is needed for normal neurodevelopment and neuron survival^52^. Notably, our results suggested differences in p75^NTR^ signaling among both neurons and glia of MDD cases and controls. While less well studied, glia also express p75^NTR^ where it is important for oligodendrocyte development and astroglial response to injury and insult^53^.

Patients with treatment-resistant MDD have reduced cortical grey matter density^54^. These grey matter reductions appear to be due to a diminished neuronal cell size paired with decreased densities of glia in MDD patients^55, 56^. Further, in rat models of depression, glial ablation in the prefrontal cortex was sufficient to induce depression-like behaviors^57^. Thus, these grey matter alterations in MDD may be partially mediated by p75^NTR^ linked apoptotic processes. While we did not observe significant differences in estimated neuron:glia ratios between MDD cases and controls in our samples, such ratios do not reflect absolute differences in cell numbers between groups.

Analysis of top bulk brain findings also demonstrated enrichment for vascular endothelial growth factor (VEGF) and nitric oxide (NO) signaling pathways. While more extensively studied as an angiogenic factor, some evidence has accumulated for VEGF as a neuroprotective factor with links to MDD^58^. In vascular endothelial cells, VEGF regulates NO synthase expression where it interacts with the p75^NTR^ in opposing fashion^59, 60^. A similar action for VEGF is seen in brain vasculature under pathological conditions where VEGF increases blood-brain barrier (BBB) permeability^61^. Thus, our findings implicating VEGF and p75^NTR^ signaling may reflect alterations in BBB integrity in MDD.

It is also interesting to note that MWAS results for neurons were significantly enriched for circadian entrainment pathways. Sleep disruption presents in 50-95% of depression cases and is correlated with severity and susceptibility to recurrent depression^62, 63^. These circadian disruptions can be seen in abnormal temporal expression of canonical clock genes in brain regions outside the suprachiasmatic nucleus, such as prefrontal cortex^64^. Whereas our MWAS results did not involve canonical clock genes, many intermediate enzymes and second messenger systems (e.g. calcium-dependent kinases, adenylyl cyclases) drove the enrichment of circadian entrainment pathways. Pathways linked to these calcium-dependent second messenger systems were also prominently featured among results for glia. Like neurons, astrocytes express circadian rhythms^65^. Given the centrality of calcium signaling in astrocyte biology^66^, glial defects may also contribute to altered circadian rhythm in MDD.

Finally, the most significant MWAS findings among blood cell types were observed in CD14+ monocytes. Considerable evidence has shown that psychological stress activates potent immune responses via the hypothalamic-pituitary-adrenal axis^67^ which leads to epigenetic reprogramming of monocytes and microglia^42, 68, 69^. These primed immune cells in-turn display a proinflammatory phenotype to future stress and strongly influence mood and behavior via neuroinflammatory processes^42, 70, 71^.

Curiously, the most significant pathway implicated in monocytes was related to glutamatergic signaling, which was also implicated in neurons. We did not observe the same methylation sites in the top of the MWAS from neurons and CD14, thereby suggesting any systemic component to MDD involves interactions between blood and brain rather than coincident changes in identical methylation sites. How glutamatergic signaling could impact monocytic biology in the context of a blood-brain interaction is not immediately clear. However, social stress has been shown to lead to depression-like behavior via disruption of the BBB72. Excess glutamate also increases BBB permeability and facilitates transmigration of monocytes into the brain^73, 74^.

While glutamate does not appear to be a chemoattractant for peripheral monocytes^75^, other glutamatergic ligands like the kynurenine metabolites kynurenic acid and quinolinic acid, can activate monocytes^75^–^77^. Further, severe depression has been associated with increased microglial production of quinolinic acid^78^, which in-turn stimulates astrocytes to secrete monocyte chemoattractant protein-1 (CCL2)79. Disruption of the BBB in depression may facilitate leakage of glutamate and/or kynurenine and its metabolites into circulation. Indeed, increased plasma concentrations of glutamate^80^ and kynurenine^81^ have been recently observed in MDD in other cohorts. Thus, methylation changes linked to glutamatergic signaling in neurons and monocytes may reflect responses to a broader excitotoxic or neuroinflammatory state. Such a model of stress-induced BBB changes and/or excitotoxicity in MDD also appear supported by our MWASs in brain involving p75^NTR^, VEGF, and cortisol pathways.

One limitation in studies involving biological samples from human patients is the potential confound of drug treatment. Since antidepressant treatment is highly correlated with MDD diagnosis, such a confound is largely unavoidable for postmortem samples. However, treatment information was available for the NESDA sample. Whereas many methylome-wide significant effects were associated with antidepressant treatment in CD3 and CD14, our top case-control findings in blood cell types did not contain any antidepressant-associated sites. Therefore, drug treatment was not a significant confound in our analyses.

In conclusion, our cell-type-specific MWASs revealed many associations otherwise obscured in bulk brain and whole blood, and provided unique mechanistic insights into the underlying disease processes. This highlights the utility of deconvolution methods as valuable approaches for performing MWAS in human samples for which only bulk tissue data is available. In addition to finding significant association signals for MDD in the neurons of the postmortem brain samples, we also found ample evidence for a role of glia in MDD. These findings are notable in light of the historically underappreciated role of glia in disease. Cell-type-specific analyses in blood strongly suggested a role for monocytes in MDD pathology and appears to corroborate animal studies linking monocytes to stress-induced behaviors, and supports a systemic model of MDD pathology. Collectively, top MWAS findings and secondary analyses pointed towards neurodegeneration and increased BBB permeability, potentially via p75^NTR^/VEGF signaling, as key components of such a systemic model. These pathways merit additional research and serious consideration as novel therapeutic targets for MDD. Lastly, top MWAS results were consistently enriched at genes previously associated with MDD, related neuropsychiatric disorders, and neurodegenerative disorders. Such overlap with external studies bolsters the veracity of our results and further highlights the shared liabilities among neuropsychiatric disorders.

## METHODS

Methods and any associated references are available in the Online Methods

## ACKNOWLEDGEMENTS

The project was supported by grant R01MH099110 from the National Institute of Mental Health. Post mortem brain tissues were received from: the Victorian Brain Bank, supported by The Florey Institute of Neuroscience and Mental Health, The Alfred and Victorian Forensic Institute of Medicine and funded by Australia’s National Health & Medical Research Council and Parkinson’s Victoria; the Stanley Medical Research Institute; The Netherlands Brain Bank, Netherlands Institute of Neuroscience, Amsterdam; the Harvard Brain Tissue Resource Center, and The Douglas – Bell Canada Brain Bank, Douglas Institute Research Center, Canada. The infrastructure for the NESDA study (www.nesda.nl) is funded through the Geestkracht program of the Netherlands Organisation for Health Research and Development (ZonMw, grant number 10-000-1002) and financial contributions by participating universities and mental health care organizations (VU University Medical Center, GGZ inGeest, Leiden University Medical Center, Leiden University, GGZ Rivierduinen, University Medical Center Groningen, University of Groningen, Lentis, GGZ Friesland, GGZ Drenthe, Rob Giel Onderzoekscentrum).

## AUTHOR CONTRIBUTIONS

RC, KA, and EvdO conceived the concept of the study and established the design. KA and EvdO executed supervision of the study. RC, JG, MZ, LX, and ZK generated the methylation data. RC, AS, JG, ZK, KA, and EvdO analyzed the data. GG, GT, BD, and BP provided expertise on biomaterial and phenotype information. RC and EvdO wrote the manuscript. All authors contributed important intellectual content to and critically reviewed the manuscript.

## COMPETING FINANCIAL INTERESTS

BP has received research funding (non-related) from Jansen Research and Boehringer Ingelheim. Other authors declare no competing financial interests.

## References

1 American Psychiatric Association. Diagnostic and Statistical Manual of Mental Disorders (American Psychiatric Association, Washington, DC, 1994).

2 Kessler, R.C., et al. The epidemiology of major depressive disorder: results from the National Comorbidity Survey Replication (NCS-R). JAMA 289, 3095–3105 (2003).

3 Depression and Other Common Mental Disorders: Global Health Estimates. (World Health Organization, Geneva, 2017).

4 Kaffman, A. & Meaney, M.J. Neurodevelopmental sequelae of postnatal maternal care in rodents: clinical and research implications of molecular insights. J Child Psychol Psychiatry 48, 224–244 (2007).

5 Szyf, M., Weaver, I.C., Champagne, F.A., Diorio, J. & Meaney, M.J. Maternal programming of steroid receptor expression and phenotype through DNA methylation in the rat. Front Neuroendocrinol. 26, 139–162 (2005).

6 Abdolmaleky, H.M., et al. Methylomics in psychiatry: Modulation of gene-environment interactions may be through DNA methylation. Am J Med Genet B Neuropsychiatr Genet 127B, 51–59 (2004).

7 Iwata, M., Ota, K.T. & Duman, R.S. The inflammasome: Pathways linking psychological stress, depression, and systemic illnesses. Brain, Behavior, and Immunity 31, 105–114 (2013).

8 Sotelo, J.L. & Nemeroff, C.B. Depression as a systemic disease. Personalized Medicine in Psychiatry 1-2, 11–25 (2017).

9 Shen-Orr, S.S. & Gaujoux, R. Computational deconvolution: extracting cell type-specific information from heterogeneous samples. Curr Opin Immunol 25, 571–578 (2013).

10 Houseman, E.A., et al. DNA methylation arrays as surrogate measures of cell mixture distribution. BMC Bioinformatics 13, 86 (2012).

11 Rahmani, E., et al. Sparse PCA corrects for cell type heterogeneity in epigenome-wide association studies. Nat Methods 13, 443–445 (2016).

12 Venet, D., Pecasse, F., Maenhaut, C. & Bersini, H. Separation of samples into their constituents using gene expression data. Bioinformatics 17 Suppl 1, S279–287 (2001).

13 Onuchic, V., et al. Epigenomic deconvolution of breast tumors reveals metabolic coupling between constituent cell types. Cell reports 17, 2075–2086 (2016).

14 Montaño, C.M., et al. Measuring cell-type specific differential methylation in human brain tissue. Genome Biology 14, R94 (2013).

15 Shen-Orr, S.S., et al. Cell type-specific gene expression differences in complex tissues. Nat Methods 7, 287–289 (2010).

16 Aberg, K.A., et al. Methylome-wide association findings for major depressive disorder overlap in blood and brain and replicate in independent brain samples. Molecular Psychiatry (2018).

17 von Bartheld, C.S., Bahney, J. & Herculano-Houzel, S. The search for true numbers of neurons and glial cells in the human brain: A review of 150 years of cell counting. Journal of Comparative Neurology 524, 3865–3895 (2016).

18 Morice-Picard, F., et al. Complete loss of function of the ubiquitin ligase HERC2 causes a severe neurodevelopmental phenotype. European Journal Of Human Genetics 25, 52 (2016).

19 Bekker-Jensen, S., et al. HERC2 coordinates ubiquitin-dependent assembly of DNA repair factors on damaged chromosomes. Nature Cell Biology 12, 80 (2009).

20 Poulsen, S.L., et al. RNF111/Arkadia is a SUMO-targeted ubiquitin ligase that facilitates the DNA damage response. The Journal of Cell Biology 201, 797–807 (2013).

21 Nagano, Y., et al. Arkadia Induces Degradation of SnoN and c-Ski to Enhance Transforming Growth Factor-β Signaling. Journal of Biological Chemistry 282, 20492–20501 (2007).

22 Schreiber, J., et al. Ubiquitin ligase TRIM3 controls hippocampal plasticity and learning by regulating synaptic γ-actin levels. The Journal of Cell Biology (2015).

23 Hung, A.Y., Sung, C.C., Brito, I.L. & Sheng, M. Degradation of Postsynaptic Scaffold GKAP and Regulation of Dendritic Spine Morphology by the TRIM3 Ubiquitin Ligase in Rat Hippocampal Neurons. PLOS ONE 5, e9842 (2010).

24 Puffenberger Erik, G., et al. A homozygous missense mutation in HERC2 associated with global developmental delay and autism spectrum disorder. Human Mutation 33, 1639–1646 (2012).

25 Xu, B., et al. Strong association of de novo copy number mutations with sporadic schizophrenia. Nature Genetics 40, 880 (2008).

26 Levy, R.J., et al. Deletion of Rapgef6, a candidate schizophrenia susceptibility gene, disrupts amygdala function in mice. Translational Psychiatry 5, e577 (2015).

27 Aberg, K.A., et al. Methylome-Wide Association Study of Schizophrenia: Identifying Blood Biomarker Signatures of Environmental Insults. JAMA psychiatry 71, 255–264 (2014).

28 Starnawska, A., et al. Hypomethylation of FAM63B in bipolar disorder patients. Clin Epigenetics 8, 52 (2016).

29 Pamela Belmonte Mahon, P.D., et al. Genome-Wide Linkage and Follow-Up Association Study of Postpartum Mood Symptoms. American Journal of Psychiatry 166, 1229–1237 (2009).

30 Roadmap Epigenomics, C., et al. Integrative analysis of 111 reference human epigenomes. Nature 518, 317 (2015).

31 Guintivano, J., Aryee, M.J. & Kaminsky, Z.A. A cell epigenotype specific model for the correction of brain cellular heterogeneity bias and its application to age, brain region and major depression. Epigenetics 8, 290–302 (2013).

32 Chan, S.L., Griffin, W.S. & Mattson, M.P. Evidence for caspase-mediated cleavage of AMPA receptor subunits in neuronal apoptosis and Alzheimer’s disease. Journal of neuroscience research 57, 315–323 (1999).

33 Sokolowski, J.D., et al. Caspase-mediated cleavage of actin and tubulin is a common feature and sensitive marker of axonal degeneration in neural development and injury. Acta Neuropathologica Communications 2, 16–16 (2014).

34 Wray, N.R., et al. Genome-wide association analyses identify 44 risk variants and refine the genetic architecture of major depression. Nature Genetics 50, 668–681 (2018).

35 Mattisson, I.Y., et al. Elevated Markers of Death Receptor-Activated Apoptosis are Associated with Increased Risk for Development of Diabetes and Cardiovascular Disease. EBioMedicine 26, 187–197 (2017).

36 Li, J., et al. Synaptic P-Rex1 signaling regulates hippocampal long-term depression and autism-like social behavior. Proc Natl Acad Sci U S A 112, E6964–6972 (2015).

37 Lin, E., et al. A Deep Learning Approach for Predicting Antidepressant Response in Major Depression Using Clinical and Genetic Biomarkers. Front Psychiatry 9, 290 (2018).

38 Penninx, B., Beekman, A. & Smit, J. The Netherlands Study of Depression and Anxiety (NESDA): Rationales, Objectives and Methods. International Journal of Methods in Psychiatric Research 17, 121–140 (2008).

39 Hattab, M.W., et al. Correcting for cell-type effects in DNA methylation studies: reference-based method outperforms latent variable approaches in empirical studies. Genome Biology 18, 24 (2017).

40 Zola, H., Swart, B., Nicholson, I. & Voss, E. Leukocyte and Stromal Cell Molecules: the CD Markers (Wiley, New Jersey, 2007).

41 Demir, S., et al. Neutrophil–lymphocyte ratio in patients with major depressive disorder undergoing no pharmacological therapy. Neuropsychiatric Disease and Treatment 11, 2253–2258 (2015).

42 Weber, M.D., Godbout, J.P. & Sheridan, J.F. Repeated Social Defeat, Neuroinflammation, and Behavior: Monocytes Carry the Signal. Neuropsychopharmacology 42, 46–61 (2017).

43 van den Oord, E.J.C.G. Controlling false discoveries in genetic studies. American Journal of Medical Genetics Part B: Neuropsychiatric Genetics 147B, 637–644 (2008).

44 van den Oord, E.J. & Sullivan, P.F. False discoveries and models for gene discovery. Trends Genet 19, 537–542 (2003).

45 Menke, A. & Binder, E.B. Epigenetic alterations in depression and antidepressant treatment. Dialogues in Clinical Neuroscience 16, 395–404 (2014).

46 Demontis, D., et al. Discovery Of The First Genome-Wide Significant Risk Loci For ADHD. bioRxiv (2017).

47 Otowa, T., et al. Meta-analysis of genome-wide association studies of anxiety disorders. Mol Psychiatry 21, 1391–1399 (2016).

48 Grove, J., et al. Common risk variants identified in autism spectrum disorder. bioRxiv (2017).

49 Ruderfer, D.M., et al. Genomic Dissection of Bipolar Disorder and Schizophrenia, Including 28 Subphenotypes. Cell 173, 1705–1715.e1716 (2018).

50 MacArthur, J., et al. The new NHGRI-EBI Catalog of published genome-wide association studies (GWAS Catalog). Nucleic Acids Research 45, D896–D901 (2017).

51 Michailidou, K., et al. Association analysis identifies 65 new breast cancer risk loci. Nature 551, 92 (2017).

52 Hempstead, B.L. The many faces of p75^NTR^. Current Opinion in Neurobiology 12, 260–267 (2002).

53 Cragnolini, A.B. & Friedman, W.J. The function of p75^NTR^ in glia. Trends in Neurosciences 31, 99–104 (2008).

54 Shah, P.J., Ebmeier, K.P., Glabus, M.F. & Goodwin, G.M. Cortical grey matter reductions associated with treatment-resistant chronic unipolar depression: Controlled magnetic resonance imaging study. British Journal of Psychiatry 172, 527–532 (1998).

55 Cotter, D., Mackay, D., Landau, S., Kerwin, R. & Everall, I. Reduced glial cell density and neuronal size in the anterior cingulate cortex in major depressive disorder. Archives of general psychiatry 58, 545–553 (2001).

56 Cotter, D., et al. Reduced Neuronal Size and Glial Cell Density in Area 9 of the Dorsolateral Prefrontal Cortex in Subjects with Major Depressive Disorder. Cerebral Cortex 12, 386–394 (2002).

57 Banasr, M. & Duman, R.S. Glial Loss in the Prefrontal Cortex Is Sufficient to Induce Depressive-like Behaviors. Biological Psychiatry 64, 863–870 (2008).

58 Clark-Raymond, A. & Halaris, A. VEGF and depression: a comprehensive assessment of clinical data. J Psychiatr Res 47, 1080–1087 (2013).

59 Shen, B.-Q., Lee, D.Y. & Zioncheck, T.F. Vascular Endothelial Growth Factor Governs Endothelial Nitric-oxide Synthase Expression via a KDR/Flk-1 Receptor and a Protein Kinase C Signaling Pathway. Journal of Biological Chemistry 274, 33057–33063 (1999).

60 Caporali, A., et al. The neurotrophin receptor p75(NTR) triggers endothelial cell apoptosis and inhibits angiogenesis: implications for diabetes-induced impairment of reparative neovascularization. Circulation research 103, e15–e26 (2008).

61 Mayhan, W.G. VEGF increases permeability of the blood-brain barrier via a nitric oxide synthase/cGMP-dependent pathway. American Journal of Physiology-Cell Physiology 276, C1148–C1153 (1999).

62 Perlis, M.L., et al. Which depressive symptoms are related to which sleep electroencephalographic variables? Biological Psychiatry 42, 904–913 (1997).

63 Wulff, K., Gatti, S., Wettstein, J.G. & Foster, R.G. Sleep and circadian rhythm disruption in psychiatric and neurodegenerative disease. Nature Reviews Neuroscience 11, 589 (2010).

64 Li, J.Z., et al. Circadian patterns of gene expression in the human brain and disruption in major depressive disorder. Proceedings of the National Academy of Sciences (2013).

65 Prolo, L.M., Takahashi, J.S. & Herzog, E.D. Circadian Rhythm Generation and Entrainment in Astrocytes. The Journal of Neuroscience 25, 404–408 (2005).

66 Perea, G. & Araque, A. Glial calcium signaling and neuron–glia communication. Cell Calcium 38, 375–382 (2005).

67 Tsigos, C. & Chrousos, G.P. Hypothalamic–pituitary–adrenal axis, neuroendocrine factors and stress. Journal of Psychosomatic Research 53, 865–871 (2002).

68 Christ, A., et al. Western Diet Triggers NLRP3-Dependent Innate Immune Reprogramming. Cell 172, 162–175.e114 (2018).

69 Fleshner, M. Stress-evoked sterile inflammation, danger associated molecular patterns (DAMPs), microbial associated molecular patterns (MAMPs) and the inflammasome. Brain, Behavior, and Immunity 27, 1–7 (2013).

70 Wohleb, E.S., Franklin, T., Iwata, M. & Duman, R.S. Integrating neuroimmune systems in the neurobiology of depression. 17, 497 (2016).

71 Wohleb, E.S., McKim, D.B., Sheridan, J.F. & Godbout, J.P. Monocyte trafficking to the brain with stress and inflammation: a novel axis of immune-to-brain communication that influences mood and behavior. Frontiers in Neuroscience 8 (2015).

72 Menard, C., et al. Social stress induces neurovascular pathology promoting depression. Nature Neuroscience 20, 1752–1760 (2017).

73 Reijerkerk, A., et al. The NR1 subunit of NMDA receptor regulates monocyte transmigration through the brain endothelial cell barrier. Journal of Neurochemistry 113, 447–453 (2010).

74 Xhima, K., Weber-Adrian, D. & Silburt, J. Glutamate Induces Blood–Brain Barrier Permeability through Activation of N-Methyl-D-Aspartate Receptors. The Journal of Neuroscience 36, 12296–12298 (2016).

75 Malone, J.D., Richards, M. & Kahn, A.J. Human peripheral monocytes express putative receptors for neuroexcitatory amino acids. Proceedings of the National Academy of Sciences of the United States of America 83, 3307–3310 (1986).

76 Tiszlavicz, Z., et al. Different inhibitory effects of kynurenic acid and a novel kynurenic acid analogue on tumour necrosis factor-α (TNF-α) production by mononuclear cells, HMGB1 production by monocytes and HNP1-3 secretion by neutrophils. Naunyn-Schmiedeberg’s Archives of Pharmacology 383, 447–455 (2011).

77 Lugo-Huitrón, R., et al. Quinolinic Acid: An Endogenous Neurotoxin with Multiple Targets. Oxidative Medicine and Cellular Longevity 2013, 104024 (2013).

78 Steiner, J., et al. Severe depression is associated with increased microglial quinolinic acid in subregions of the anterior cingulate gyrus: Evidence for an immune-modulated glutamatergic neurotransmission? Journal of Neuroinflammation 8, 94 (2011).

79 Guillemin, G.J., Croitoru-Lamoury, J., Dormont, D., Armati, P.J. & Brew, B.J. Quinolinic acid upregulates chemokine production and chemokine receptor expression in astrocytes. Glia 41, 371–381 (2003).

80 Inoshita, M., et al. Elevated peripheral blood glutamate levels in major depressive disorder. Neuropsychiatric Disease and Treatment 14, 945–953 (2018).

81 Pan, J.-X., et al. Diagnosis of major depressive disorder based on changes in multiple plasma neurotransmitters: a targeted metabolomics study. Translational Psychiatry 8, 130 (2018).

